# Multiple geographic breakdown events of the *Rpv1*–*Rpv3.1* pyramided resistance in grapevine by *Plasmopara viticola*

**DOI:** 10.1101/2025.08.25.672106

**Authors:** Rémi Pélissier, François Delmotte, Laurent Delière, Julie Ramirez Martinez, Laura Marolleau, Isabelle D. Mazet, Frédéric Fabre, Anne-Sophie Miclot

**Affiliations:** INRAE, Bordeaux Sciences Agro, SAVE, ISVV, F-33140 Villenave d’Ornon, France; INRAE, Bordeaux Sciences Agro, SAVE, UE Vigne Bordeaux, ISVV, F-3882, Villenave d’Ornon, France

**Keywords:** *Vitis vinifera*, grapevine downy mildew, disease resistance, convergent evolution, gene pyramiding, Resistance breakdown

## Abstract

In 2024, three vineyard sites located within a radius of 40 km in southeastern France, each planted with Artaban, a grapevine variety carrying pyramided resistance (*Rpv1* and *Rpv3.1*), exhibited severe downy mildew symptoms following climatic conditions highly conducive to disease development. Twenty-one *Plasmopara viticola* isolates were collected from these vineyards and, after monosporangium isolation, tested under controlled conditions on a panel of susceptible and resistant grapevine varieties to assess their virulence. Among the 21 tested field strains, 19 overcame *Rpv3.1* mediated resistance, 10 overcame *Rpv1* alone, and four genetically distinct strains from two separate vineyards were overcoming the pyramided *Rpv1*–*Rpv3.1* combination. The genetic characterization of the strains suggests that this adaptation likely resulted from convergent evolution. To our knowledge, these represent the first documented cases in Europe of a breakdown in resistance conferred by *Rpv1*, whether deployed alone or in combination with *Rpv3.1*.

## Introduction

Grapevine downy mildew (GDM), caused by the oomycete pathogen *Plasmopara viticola*, is one of the most destructive diseases worldwide (Millardet, 1881), making grapevine an intensively treated crops, as its control still depends largely on fungicide applications (Gessler *et al*., 2011; Koledenkova *et al*., 2022; Nefti *et al*., 2024). An ongoing sustainable alternative involves breeding and deploying disease-resistant varieties (DRVs). Several resistance factors, originating from wild *Vitis* species have already been identified and are currently being incorporated into breeding programs (Merdinoglu *et al*., 2018; Possamai & Wiedemann-Merdinoglu, 2022). Four of them (*Rpv1*, *Rpv3.1*, *Rpv10*, and *Rpv12*) are currently the most widely used in DRVs. However, as resistance genes are a limited resource and can be overcome by pathogens, their deployment should be carefully managed to prevent, as much as possible, the emergence and fast spread of resistance-breaking strains (Parlevliet, 2002; Merdinoglu *et al*., 2018; Rimbaud *et al*., 2021).

Resistance breakdowns have already been observed in *P. viticola* populations despite the limited deployment of DRVs. For instance, the resistance factor *Rpv3.1*, one of the resistance most deployed in Europe, is today largely overcome (Peressotti *et al*., 2010; Delmotte *et al*., 2014; Delmas *et al*., 2016; Wingerter *et al*., 2021; Paineau *et al*., 2022). The first resistance breakdown was detected in 2010 in Czech Republic, in the Bianca variety, which carries the *Rpv3.1*, one of the 3 haplotypes (genetic variants of the resistance locus) of the *Rpv3* factor (Peressotti *et al*., 2010). This widespread adaptation to Rpv3.1 observed in contemporary GDM populations is likely the result of the large deployment of French-American hybrids during the first half of the 20th century (Di Gaspero *et al*., 2012). Recently, several *P. viticola* strains exhibiting high aggressiveness on varieties carrying the *Rpv10* factor (Delmas *et al*., 2016; Heyman *et al*., 2021) or even overcoming this resistance (Paineau *et al*., 2022) have been reported. Similarly, three strains breaking down *Rpv12* factor have been isolated in Hungary, Germany and Italy (Wingerter *et al*., 2021; Paineau *et al*., 2022; Dvorak *et al*., 2025). The genomic determinants of *P. viticola* virulence against these resistance factors have been documented and largely conform to a gene-for-gene model. Avirulence loci associated with *Rpv3.1* and *Rpv12* containing multiple several candidate RXLR effector genes have been identified (Paineau *et al*., 2024; Dvorak *et al*., 2025). Virulent strains overcoming *Rpv3.1* and *Rpv12* carry large homozygous deletions encompassing several RXLR genes within these loci. In the case of *Rpv10*, virulence was found to be dominant and associated with an admixed genomic segment likely originating from a secondary introduction of GDM into Europe (Dvorak *et al*., 2025).

By contrast, the *Rpv1* factor, originated from *Muscadinia rotundifolia* (Merdinoglu *et al*., 2003), remain the only one that has not yet been overcome in Europe. This factor, which confers quantitative (partial) resistance against GDM, is a NB-LRR receptor, with a nucleocytoplasmic location, involved in pathogen recognition and signal transduction during the initiation of plant defense (Feechan *et al*., 2013). To date, only one strain overcoming this resistance has been isolated on Réunion Island in the Indian Ocean (Ramirez *et al*. in prep), but the effector in GDM recognize by this receptor remains unknown. *Rpv1* has been used in breeding programs developed in France since the 1980s to create a series of monogenic DRV called Bouquet varieties (Salmon *et al*., 2018). They were subsequently used in the breeding program ResDur developed by INRAE and IFV since the early 2000s to obtain pyramided (i.e. polygenic) DRV. These varieties combine the resistance genes *Rpv1* and *Rpv3.1* originating from german varieties. A first set of four pyramided ResDur varieties (Artaban, Floreal, Vidoc and Voltis) have been registered in the French official cultivar catalogue in 2018. Bouquet varieties were also used to combine *Rpv1* with *Rpv10*, a gene originating from Asian *Vitis* species introgressed through the Bronner variety. This effort results in the registration in 2021 of eight ResDur varieties of second-generation (Coliris, Lilaro, Sirano, Selenor, Calys, Artys, Exelys and Opalor) (Yobrégat, 2018; Schneider *et al*., 2019). Currently, *Rpv1* remain a central component in ongoing breeding programs, including the development of the third generation of ResDur varieties (combining *Rpv1, Rpv3.1 and Rpv10)* particularly because of its linkage with the resistance gene *Run1*, which provides complete resistance to powdery mildew (Merdinoglu *et al*., 2003; Agurto *et al*., 2017).

The 2024 cropping season was marked by a severe GDM epidemic affecting vineyards across France. For the first time, three fields in southeastern France planted with the Artaban variety, carrying the pyramided *Rpv1*-*Rpv3.1* factor, exhibited high downy mildew severity on both clusters and leaves. To test the hypothesis of a possible adaptation of the grapevine downy mildew population to the resistant variety Artaban in these fields, we collected 21 *Plasmopara viticola* isolates and performed both genetic and phenotypic characterization. The virulence profiles of these isolates were assessed through cross-inoculation experiments under controlled conditions, evaluating both sporulation ratio and hypersensitive response (HR) across the main resistance genes currently deployed in Europe. Genetic characterization was conducted using 10 SSR markers, enabling to describe the genetic structure of this population of strains.

## Material and methods

### Disease field notation and pathogen collected

GDM epidemics were assessed in July 2024 from three fields planted with the DRV Artaban. These vineyards (or field) were located in the Rhône Valley wine-growing area (South-east, France). These fields were conducted without fungicide treatments.

Disease in the field was assessed on both leaves and grape clusters using a standardized scoring scale developed for the OSCAR observatory (Guimier *et al*., 2019). We recorded, for each field, the incidence of infected plants that is the proportion of plants exhibiting visible symptoms, separately for leaves and clusters. This continuous value is then transformed into ordinal classes, as follows: 0: No disease detected; 1: Trace of disease (<5%); 2: symptoms visible regularly (5–25%); 3: important presence (25–50%); 4: Very important presence (50– 80%); 5: symptoms always present (80–100%). In addition, we quantified symptom severity by estimating the average percentage of tissue affected on symptomatic plants, a value subsequently transformed into ordinal classes as follow: 0: No disease detected; 1: Trace of disease (<1%); 2: Clearly visible symptoms (1–5%); 3: Significant damage (5–10%); 4: Very severe damage (10–50%); 5: Extremely severe damage (>50%).

### Pathogen isolation and multiplication

In total, 21 downy mildew isolates, 11 from Caderousse, 6 from Ancone and 4 from St-Geniès-de-Malgoirès were sampled from infected plants **(Figure 1)**. In addition, 8 reference strains with known virulence profiles were inoculated: 5 avirulent strains (REF *Avir*), 2 virulent on *Rpv*3.1 (REF vir3.1) and 1 virulent on *Rpv*3.1 and *Rpv*12 (REF vir3.1, 12) (**Table S1**).

**Figure 1.**
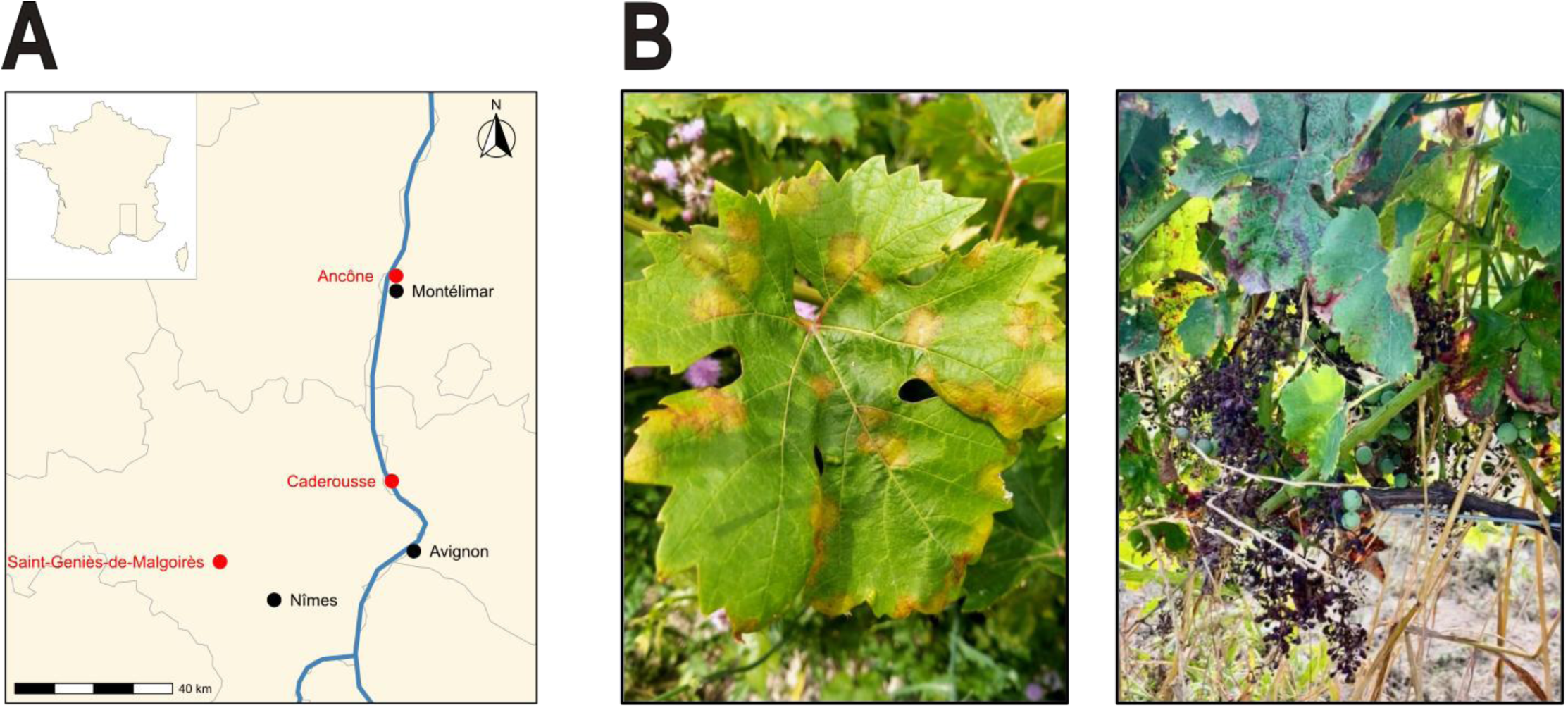
Localization and type of symptoms observed on Artaban in the three sampled vineyards **A)** Map showing the locations of the sampled vineyards (Caderousse, Ancône, and Saint-Geniès-de-Malgoirès, indicated in red), where samples were collected from Artaban (*Rpv1*-*Rpv3.1*) fields severely affected by downy mildew in 2024. Names of the main nearby cities are shown in black for reference. **B)** Symptoms of downy mildew observed on grapevine leaves and clusters in the field in Caderousse.

Each isolate consisted of a single sporulating lesion collected from a single infected leaf. The leaf fragments were rinsed with sterile water and left overnight in the dark to obtain sporulation. Fresh sporangia were collected and stored in liquid nitrogen after desiccation for subsequent experiments. For each isolate, the sporulating leaf fragments stored in liquid nitrogen were gently agitated against a microscope slide to release the sporangia. Under a binocular microscope, a single sporangium was caught with a disinfected human eyelash and gently deposited on a 15 μl droplet of reverse-osmosis water at the center of a 15-mm-diameter leaf disc cut from a V. vinifera ‘Cabernet Sauvignon’ plant. The inoculated discs were placed overnight in the dark. The water droplets were removed by suction and then the discs were incubated for 6 days at 22°C, under a 12-h light/12-h dark photoperiod. The infection efficiency of a single sporangium is low (about 10%). We therefore isolated 10 sporangia for each isolate in this way. After 6 days of incubation in a growth chamber, the infected leaf discs (one disc per isolate) were placed in Eppendorf tubes and left overnight in a desiccator before storage at −20°C. The individuals obtained by monosporangium isolation are referred to hereafter as strains. Two weeks before the experiment, the strains were propagated on five different leaf fragments of Cabernet Sauvignon. After 1 week of incubation, they were then propagated on four detached leaves for the cross-inoculation experiment. One day before the experiment, the sporulating leaves were gently rinsed with distilled water to remove the sporangia already present, to ensure the collection of fresh sporangia.

### Plant material and cross-inoculation experiments

All grapevine plants were obtained from commercial nurseries and grafted onto SO4 rootstock. All grapevine varieties were grown simultaneously in a greenhouse under natural photoperiod conditions, without chemical treatment. Cross-inoculation experiments were realized on leaf discs of two susceptible varieties (Chardonnay and Cabernet Sauvignon) and 7 resistant varieties (Coutia carrying *Rpv1*; Regent carrying *Rpv3.1*; Artaban and Floreal carrying *Rpv1*-*Rpv3.1 and Fleurtai carrying Rpv12*). The cross-inoculation experiment was conducted on leaves collected after 6 weeks of cultivation. Inoculations were performed on the fifth leaf below the apex. Leaves were washed with distilled water and dried on absorbent paper. We excised leaf discs with a diameter of 15 mm with a cork borer and placed them, abaxial side up, on wet filter paper in a Petri dish. The four-leaf discs inoculated for a given interaction “Strains*Inoculated variety”, were collected from different plants.

Sporangial suspensions of *P. viticola* were prepared in sterile water maintained below 10 °C, and adjusted to a concentration of 20,000 sporangia/mL using a portable particle counter (Scepter 2.0 automated cell counter; Millipore). Suspensions were kept on ice until use. Inoculation was carried out by depositing a 50 µL drop of the suspension onto the upper surface of each leaf disc. For each *P. viticola* strain, we inoculated one Petri dish containing 4 leaf discs of each variety. Mock-inoculated controls, consisting of sterile water only, were included in each experimental setup. One day post-inoculation (DPI 1), residual water droplets were removed by gently blotting the surface of each Petri dish using single-use absorbent paper. Immediately afterward, Petri dishes were sealed with Parafilm to ensure 100% relative humidity. Inoculated samples were then incubated for five days at 22 °C under a 12-hour light/12-hour dark photoperiod.

### Phenotypic characterization of the strains, and statistical analysis

Five days post-inoculation, all leaf discs were photographed on the upper and lower sides. The sporulation ratio was determined by image analysis on the lower side of each disc, as the ratio between the sporulating surface of the disc and its total surface. The code used is available as a Jupyter Notebook at https://gitlab.com/grapevinedownymildew/notebook_image_analysis.

The droplet inoculation method induces sporulation only on a localized area corresponding to the size of the droplet, approximately one tenth of the total leaf disc surface. As a result, the sporulation ratios estimated in this study need to be multiplied by a factor of approximately 10 to align with values reported in studies using the spray inoculation method as performed in Paineau et al 2022 (which enables sporulation over the entire leaf disc).

Leaf necrosis was determined visually using a categorical score characterizing the type of necrosis adapted from Paineau et al. 2022. Specifically, the type of necrosis was assessed on the upper side of the discs based on the shape, color, and size of necrosis as qualitative variable (**Figure S1**). Only necrosis types from 2 to 5 were considered as hypersensitive response (HR). Importantly, these scores are not ordinal; they describe the type or pattern of the necrosis, rather than the intensity of necrosis.

The definition of resistance breakdown was adapted from Paineau et al. 2022. The virulence of a strain on a given *Rpv* factor require the absence of a hypersensitive response by the plant and the occurrence of sporulation on both the susceptible and the resistant plants (**Figure 2**). These two conditions allow the identification of strains overcoming (i.e. breaking down) the considered resistance factor. By contrast, strain with at least one disc exhibiting HR were classified as avirulent.

**Figure 2.**
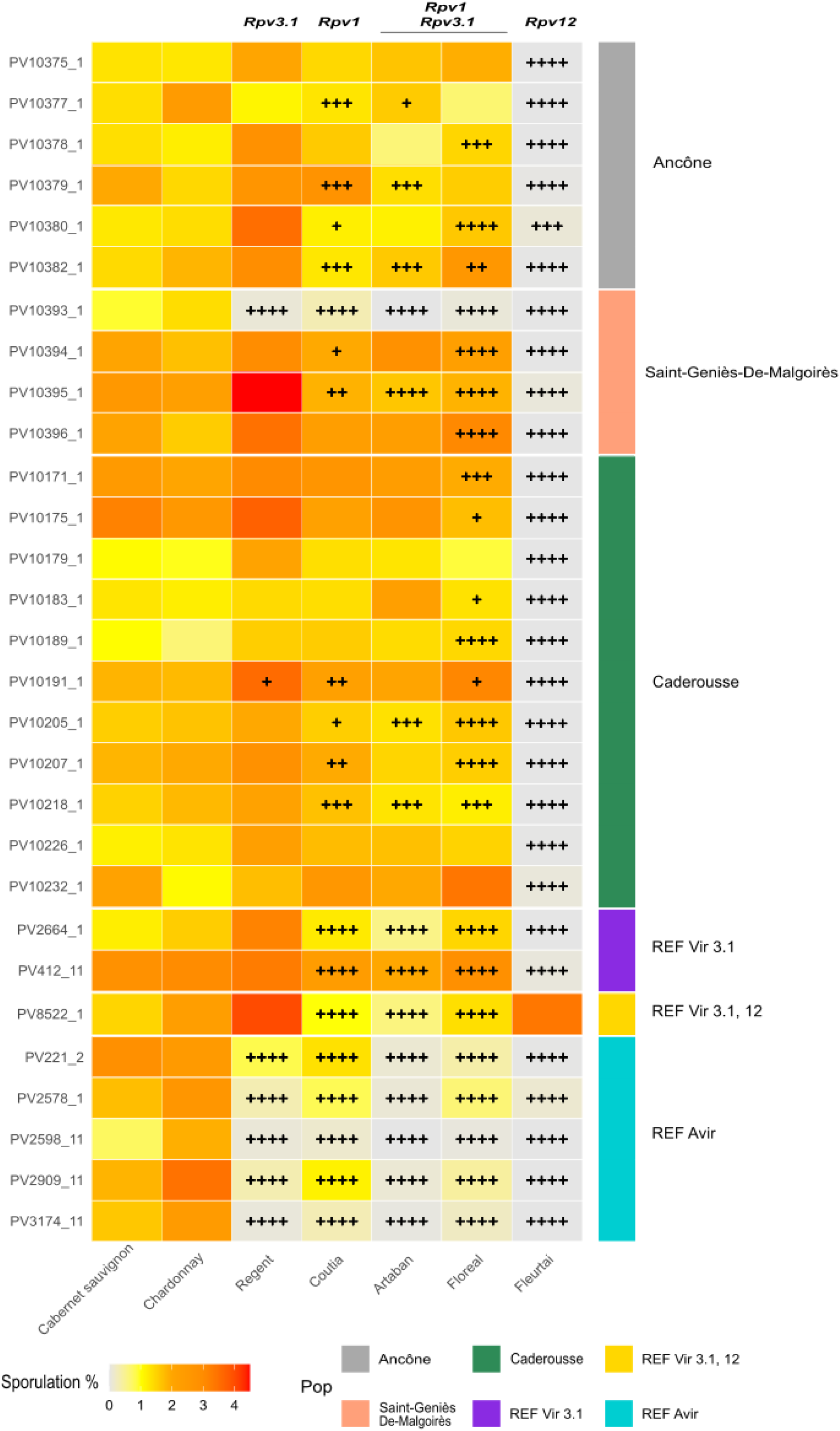
Sporulation ratio mean and leaf Hypersensitive Response (HR) of the 21 downy mildew strains collected in the field and the 8 reference strains.

For the statistical analyses, we first tested whether the sporulation ratios depend on the virulence profile of the strains for each inoculated variety. The sporulation ratios were square root transformed to meet the assumption of normality. The virulence profile is defined by a qualitative variable with 3 levels: REF Avir (5 strains), REF vir3.1 (2 strains), and vir1,3.1 (the four field-derived resistance-breaking strains). A linear mixed-effects model was used to analyze the impact of the virulence profile on the sporulation ratios. The model included the virulence profile as a fixed effect and a random intercept for individual strain nested within virulence profile to account for the non-independence of measurements taken from isolates belonging to the same group (**Figure 3**).

**Figure 3.**
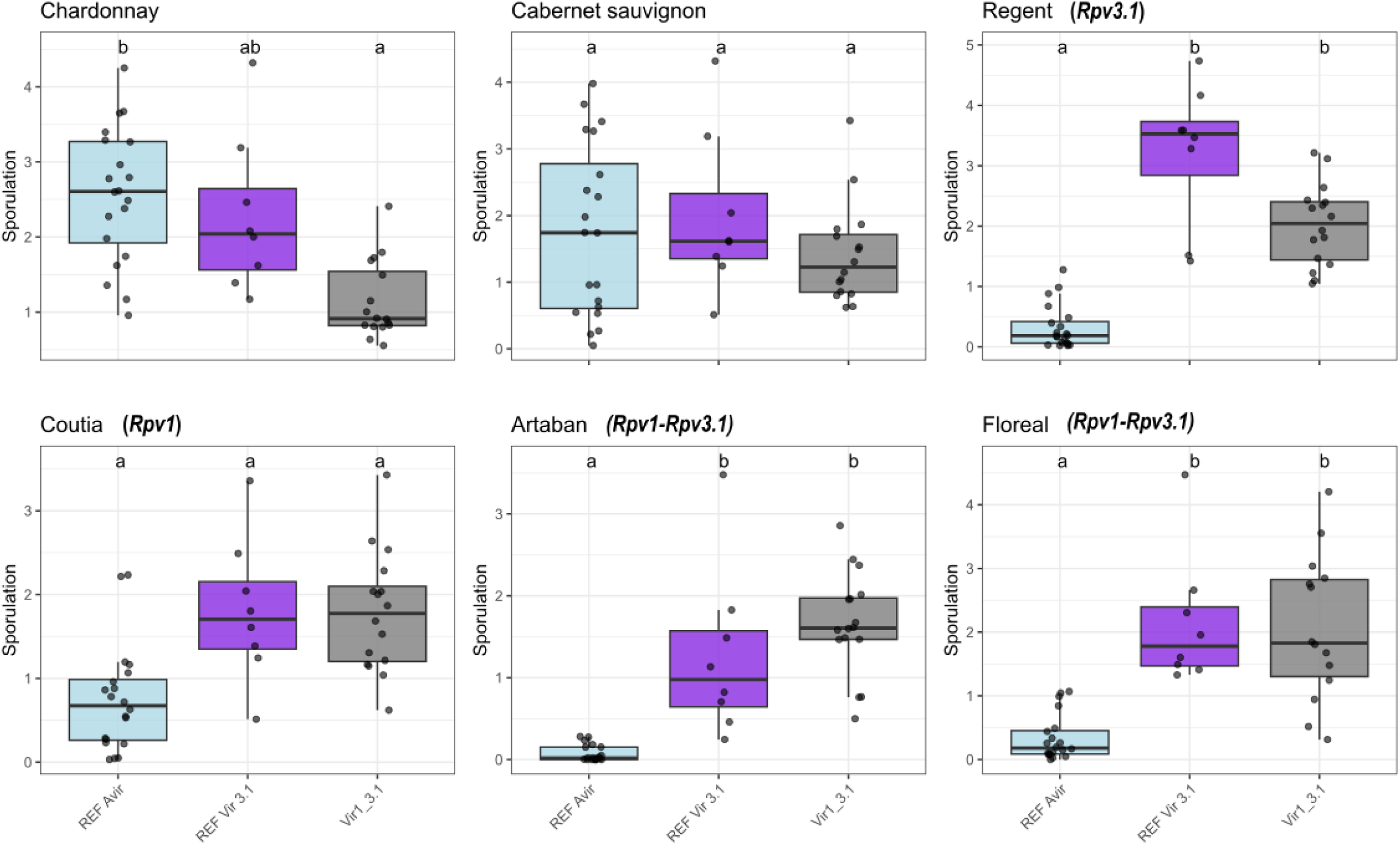
Comparison of sporulation ratio of the strains overcoming *Rpv1* and *Rpv3.1* and the reference strains. Boxplots show the sporulation ratio (as a percentage of infected leaf disc surface) for four field strains identifed as *Rpv1*-*Rpv3.1* breaking strain (Vir1,3.1: PV10226_1, PV10232_1, PV10375_1, PV10179_1), avirulent reference strains (REF Avir: PV221_2, PV2578_1, PV2598_11, PV2909_11, PV3174_11), and virulent reference on *Rpv3.1*(REF Vir3.1: PV2664_1,PV412_11) inoculated on six grapevine varieties: Chardonnay and Cabernet Sauvignon (susceptible), Coutia (*Rpv1*), Regent (*Rpv3.1*), Artaban and Floreal (*Rpv1-Rpv3.1*). Sporulation ratio was assessed via image analysis. Letters above the boxplots indicate significant groupings based on Tukey’s HSD post-hoc test (α = 0.05). Colors denote the strain origin: blue for avirulent, purple and red for virulent references, and grey for the tested field strains

Secondly, we grouped the strains into two categories, the four field strains adapted to the *Rpv1* and *Rpv3.1* on one side, and the 15 field strains only adapted to the *Rpv3.1* locus on the other side. For each category, we tested whether the sporulation ratios depend on the qualitative variable inoculated variety (6 levels) using a linear model and a one-way ANOVA (**Figure S4**). Thirdly, we tested whether the sporulation ratios depend on the inoculated variety (6 levels), the level of adaptation of the strains to the *Rpv1* factor and their interaction using a linear model with interaction. The adaptation level of the strains to the *Rpv1* factor is a quantitative variable defined as the number of necrotic leaf discs (score ≥ 2) observed on variety with *Rpv1* (Artaban, Coutia, Floreal; **Figures S3 and S5**).

For each analysis, normality and homoscedasticity were verified by inspecting the residuals. When appropriate, Tukey’s HSD post-hoc tests were used to identify significant differences between the levels of the qualitative variables considered.

All statistical analyses and data visualizations were performed using R (version 2024.12.0; R Core Team, 2024). For statistic, the following R packages were used:, ggplot2 for data visualization (Wickham, 2016), lme4 for fitting linear mixed-effects models (Bates *et al*., 2015), emmeans for estimated marginal means and post hoc comparisons (Lenth, 2017), multcomp for multiple comparisons (Bretz *et al*., 2008). The analyses can be reproduced using the Rmd file provided as supplementary material.

### Genetic analysis

The DNA from 3 reference strains, i.e. PV2909_1 (REF Avir), PV2664_1 (REF vir3.1), PV8522_1 (REF vir3.1,12) and from 21 field strains collected in the 3 fields of Artaban was extracted directly from the mycelium of *P. viticola* as described by Dussert *et al*. (2019) using a CTAB-based protocol adapted from Möller *et al*. (1992).

To assess the genetic structure of *P. viticola* strains collected from the fields, we genotyped each strain using 10 previously published microsatellite markers (isa, Pv13, Pv14, Pv16, Pv137, Pv140, Pv141, Pv142, Pv143, Pv144) and performed the analysis as described in Delmotte *et al*., 2006 and Rouxel *et al*., 2012. The Bruvo’s distance (Bruvo *et al*., 2004) was calculated on the multilocus genotype data using the poppr package (Kamvar *et al*., 2014), with repeat lengths set uniformly to two for all loci. A NJ tree was then constructed from the Bruvo distance matrix using the ape package (Paradis *et al*., 2004). The resulting unrooted tree was visualized using the ggtree package (Yu *et al*., 2017). Node labels were annotated with strain names, and tips were colored according to their geographic origin. A Bruvo’s genetic distance threshold of 0.05 was used to group multilocus genotypes into clonal lineages, accounting for minor allelic differences potentially due to somatic mutations.

## Results

### Field disease notation

We evaluated GDM incidence and severity in the three fields inspected. Foliar incidence was consistently high, with more than 80% of plants infected across all fields. Symptom severity on leaves was also high, exceeding 50% in two locations (Caderousse and Ancône) and lower, between 5% and 10%, in the third location Saint-Geniès-de-Malgoirès. Similarly, disease incidence on clusters was over 80% across all three sites. Disease severity on clusters was particularly severe in Caderousse (>50%), while it ranged between 10% and 50% in Ancône and Saint-Geniès-de-Malgoirès (**Table 1**, **Figure 1**).

**Table 1:** Field Characteristics and disease scoring of the three vineyards sampled DRV: Disease Resistant Variety; GDM: Grapevine Downy Mildew.

| Location | Grapevine DRV | Years of plantation | Field area | GDM Leaf incidence | GDM Leaf severity | GDM cluster incidence | GDM cluster severity |
| --- | --- | --- | --- | --- | --- | --- | --- |
| Caderousse | Artaban | 2021 | 4 ha | 5/5 (>80%) | 5/5 (>50%) | 5/5 (>80%) | 5/5 (>50%) |
| Ancône | Artaban | 2020 | 2,2 ha | 5/5 (>80%) | 5/5 (>50%) | 5/5 (>80%) | 4/5 (10-50%) |
| Saint Geniès-de-Malgoirès | Artaban | 2020 | 3 ha | 5/5 (>80%) | 3/5 (5-10%) | 5/5 (>80%) | 4/5 (10-50%) |

Sporulation ratio and HR were assessed for each field-collected strains and reference strains on 4 leaf discs from susceptible varieties (Chardonnay and Cabernet Sauvignon) and resistant varieties carrying different *Rpv* factor: Regent (*Rpv3.1*), Coutia (*Rpv1*), Artaban and Floreal (*Rpv1-Rpv3.1*) and Fleurtai (*Rpv12*). The mean of sporulation ratio is represented using a color scale from grey (no sporulation) to red (more than 5% of the disc surface sporulating). A “+” symbol indicates a disc (out of 4) presenting a necrosis type ≥ 2, interpreted as HR and evidence of avirulence on the corresponding variety (detail is available on Figure S2). Field strains are color-coded according to their geographic origin.

### Strain virulence characterization

We inoculated 21 strains isolated on Artaban (*Rpv1*-*Rpv3.1*) (11 from Caderousse, 6 from Ancône and 4 from Saint-Geniès-de-Malgoirès), and 8 reference strains comprising 5 avirulent strains and 3 strains having different virulence profiles (**Table S1**). Field strains virulence was determined based on the type of necrosis (if any) and the occurrence of sporulation (**Figure 2**).

### Hypersensitive response

Virulence was first assessed based on the type of necrosis. Strains with at least one disc exhibiting HR of (necrosis type 2, 3, 4, or 5) were considered as avirulent. By contrast, strains with all discs showing sporulation and scored as necrosis type 0 or 1 were classified as virulent.

Reference avirulent strains induced HR without or with low sporulation on each of the *Rpv* factors tested. Virulent reference strains vir3.1 (PV2664_1 and PV412_11) sporulated without inducing HR on Regent (*Rpv3.1*). Consistent results were obtained on Fleurtai (*Rpv12*), where the reference strain PV8522_1 (vir3.1,12) sporulated without triggering HR. On varieties carrying *Rpv1* alone or in combination with *Rpv3.1*, both vir3.1 and vir3.1,12 strains exhibited a hypersensitive response (HR), indicating their avirulence on this resistance factor.

Among the 21 strains, only two (PV10393_1 and PV10191_1) triggered an HR on the Regent (*Rpv3.1*). The other 19 strains sporulated without inducing HR and were classified as virulent (i.e. breaking down) on *Rpv3.1* **(Figure 2; Figure S2)**. From this set of 19 strains, we assessed the number of leaf discs showing HR on varieties carrying *Rpv1* alone (Coutia) or in combination with *Rpv3.1* (Artaban and Floreal). Ten out of 19 strains were virulent on Coutia (*Rpv1*), 13 were virulent on Artaban (*Rpv1*-*Rpv3.1*) and 6 were virulent on Floreal (*Rpv1*-*Rpv3.1*).

Importantly, 4 of these 19 strains (PV10232_1, PV10226_1, PV10179_1, and PV10375_1) sporulated without inducing HR on Coutia (*Rpv1*), Artaban or Floreal (*Rpv1*-*Rpv3.1*), (0/12 discs; **Figure 2; Figure S2-3**). These strains were thus breaking down the resistance conferred by both *Rpv1* and *Rpv3.1*. Importantly, they were sampled in two locations, 3 strains from Caderousse and 1 strain from Ancône, 55 km apart.

### Sporulation ratio

For each strain, sporulation ratios were interpreted relative to the levels observed on susceptible varieties and compared to the levels obtained with reference strains.

The pool of 19 strains overcoming the *Rpv3.1* factor showed higher sporulation on Regent (*Rpv3.1*) than on the susceptible variety Chardonnay and Cabernet Sauvignon, similarly to what was observed for reference strains. This same pattern was also observed for the 4 strains overcoming the combination *Rpv1* and *Rpv3.1*. These strains were sporulating significantly more on the DRV Regent (*Rpv3.1*) and Floreal (*Rpv1*-*Rpv3.1*), and tended to sporulate more on Coutia (*Rpv1*) and Artaban (*Rpv1*-*Rpv3.1*), compared to their sporulation ratio on susceptible varieties (**Figure S4**).

Compared to the pool of 5 avirulent reference strains, the 4 field strains overcoming the combination *Rpv1* and *Rpv3.1* showed significantly higher sporulation ratios on Regent (*Rpv3.1*), Artaban and Floreal (*Rpv1-Rpv3.1*) but not on Coutia (*Rpv1*). These patterns are consistent with those observed for the reference strains vir3.1. Interestingly, the 4 field strains sporulated at levels comparable to avirulent reference strains on the susceptible variety Cabernet Sauvignon but significantly less on Chardonnay (**Figure 3**).

We analyzed the relationship between the number of discs showing HR on varieties carrying *Rpv1*, and the sporulation ratio. The number of discs showing HR is considered as a proxy of the adaptation level of the strains to the *Rpv1* factor (**Figure S3-5**). The number of discs showing HR does not affect the sporulation ratio of the strains on the DRV Coutia (*Rpv1*), Artaban (*Rpv1-Rpv3.1*), and Floreal (*Rpv1-Rpv3.1*) (**Figure S5**). By contrast, the sporulation ratio significantly increases with the frequency of HR induction for the DRV Regent (*Rpv3.1*), showing that strains adapted to only *Rpv3.1* sporulate more on Regent (*Rpv3.1*) than strains adapted to *Rpv3.1* and *Rpv1* together. The same significant interaction was also found on the susceptible variety Chardonnay, suggesting that strains less adapted to *Rpv1* have higher sporulation ratio on this variety.

Taken together, the analysis of the HR type and the sporulation ratio suggest that, out of the 21 tested strains, one is avirulent (PV10393_1), 19 overcome *Rpv3.1* in Regent, 10 overcome *Rpv1* alone in Coutia (*Rpv1*) and 4 fully overcome the resistance conferred by *Rpv1* alone, *Rpv3.1* alone, and by their combination in Artaban and Floreal (*Rpv1*-*Rpv3.1*). These strains exhibited higher sporulation ratios on resistant varieties (except Coutia) than on susceptible ones. Among these 4 strains, 3 originated from the Caderousse vineyard and 1 from Ancône.

### Strain Clonality and Genetic Structure

To identify putative genetic relationships among the *P. viticola* strains, we genotyped the 21 field strains (1 sample failed amplification for PV10379_1) and 3 reference strains PV2909_1 (REF Avir), PV2664_1 (REF Vir3.1), PV8522_1 (REF Vir3.1,12).

A total of 17 multilocus genotypes (MLG) were detected among the 20 strains field strains amplified. Clonal groups were identified, with 3 of these MLGs being shared by 2 to 4 strains (using a Bruvo’s genetic distance threshold of 0.05 to define clonal groups). For instance, the strains PV10375_1, PV10378_1, PV10380_1 and PV10382_1 which were collected in the same field in Ancône shared identical genotypes across all microsatellite loci, strongly supporting a clonal propagation of the same individual. Similarly, two other clonal group were detected, one in the Saint-Geniès-De-Malgoirès field (PV10396_1, PV10394_1) and one in the Caderousse field (PV10171_1, PV10175). However, the four strains overcoming the resistance conferred by *Rpv1* alone, *Rpv3.1* alone, and by their combination display different multilocus genotypes. Additionally, no identical genotypes were identified between strains originating from different vineyards. (**Figure 4**).

**Figure 4.**
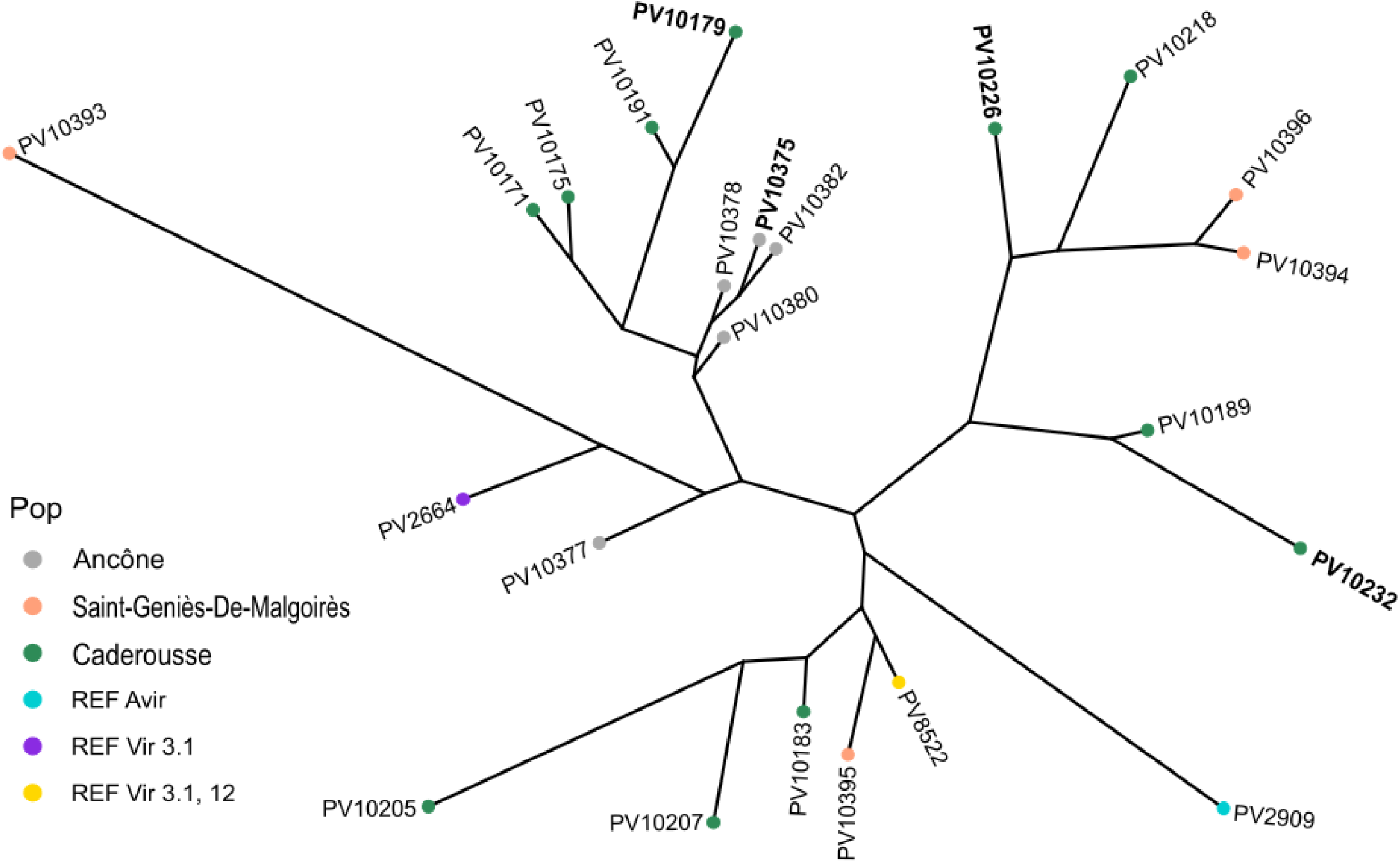
Phylogenetic analysis of the field and reference strains. Neighbor Joining unrooted tree based on allele Bruvo distance calculated with ten microsatellite loci for three reference strains of this study PV2909 (*REF Avir*), PV2664 (*REF Vir3.1*), PV8522 (*REF Vir3.1,12*) and 20 strains collected on Artaban (*Rpv1*-*Rpv3.1*) variety in Ancône, Saint-Geniès-de-Malgoirès and Caderousse fields. Each strain is color-coded according to its geographic origin. Field strains identified as *Rpv1-Rpv3.1* breaking strains (*vir1,3.1*) are written in bold.

We further investigated whether the population of strains conformed to Hardy-Weinberg expectations. Among the 10 loci analyzed, 7 (Pv14, Pv16, Pv137, Pv141, Pv142, Pv143, Pv144) were in Hardy-Weinberg equilibrium (p > 0.05), indicating that the population is broadly consistent with panmixia.

## Discussion

Pyramided resistant grapevine varieties have proven effective in controlling grapevine downy mildew on leaves and clusters in natural conditions (Calonnec *et al*., 2013; Miclot *et al*., 2022), and are expected to be a major tool for the durable management of disease resistance (Merdinoglu *et al*., 2018; Bove *et al*., 2019; Rimbaud *et al*., 2021). However, for the first time in Europe our study identifies and characterizes four *P. viticola* strains sampled in two separate vineyards, overcoming the resistance conferred by the combination of *Rpv1* and *Rpv3.1.* These strains exhibited higher sporulation levels than avirulent strains across resistant varieties carrying the pyramided resistance, without triggering any hypersensitive response, a breakdown phenotype already demonstrated for other *Rpv* factors (Delmotte *et al*., 2014; Paineau *et al*., 2022). To our knowledge, this study reports the first breakdown of *Rpv1,* alone or in combination with *Rpv3.1* in Europe. This point is especially significant as *Rpv1* was the last resistance factor in commercially available grapevine cultivars that had not been overcome by *P. viticola* in Europe. On a broader scale, nearly all strains collected from the Artaban (*Rpv1*-*Rpv3.1*) fields are characterized by a low frequency of HR by DRVs carrying *Rpv1,* compared to avirulent reference strains, suggesting that the *Rpv1* resistance factor only partially recognizes these strains. Interestingly, differences in HR frequency were observed between Artaban and Floreal (*Rpv1*-*Rpv3.1*), despite both genotypes sharing the same *Rpv* factor. As previously shown for *Rpv3.1* (Foria *et al*., 2018), such variation may result from differences in their genetic background, shaped by their respective pedigrees and process of *Rpv1* introgression. Although Coutia (*Rpv1*), Artaban and Floreal (*Rpv1*-*Rpv3.1*) descend from a common grandparental line, Artaban shares a direct parent with Coutia (Salmon *et al*., 2018). This closer relatedness may explain the phenotypic similarity in number of strains triggering HR between Artaban (*Rpv1*-*Rpv3.1*) and Coutia (*Rpv1*), in contrast to what is observed in Floreal (*Rpv1*-*Rpv3.1*).

In 2024, weather conditions strongly favored the development of downy mildew in France, in particular with significantly above-average rainfall totals in several wine growing regions. Whereas the south-east region where the samples were collected is typically not conducive to downy mildew development (Thind *et al*., 2004; Delière *et al*., 2015), the high rainfall (21% above average) and temperature (1.2°C higher than normal) in 2024 (‘Bilan climatique 2024 en France | Météo-France’) strongly favor a severe downy mildew epidemic. These favorable climatic conditions, combined with the absence of fungicide applications in the Artaban fields, contributed to high pathogen population sizes thereby facilitating the emergence of virulent strains (McDonald & Linde, 2002). Genetic analysis reveals that the *vir1* strains collected in Caderousse and Ancône are genetically distinct, suggesting independent emergences of virulent strains in each location, likely through convergent evolution. We identified a few strains sharing the same multilocus genotype within the same fields, an observation consistent with the local clonal reproduction. However, the vast majority of strains are genetically distinct, highlighting the contribution of sexual reproduction in the demo-genetic dynamics of *P. viticola* epidemics, as previously reported (Rouxel *et al*., 2012; Fontaine *et al*., 2021). The fact that the three vir1 strains from the Caderousse field are genetically different individuals suggests either three independent mutation events or, more likely, the recombination of the virulence trait into three genotypes during previous annual sexual reproduction cycles. Although highly concerning, these events remain rare in view of the more than 1,000 hectares currently planted with ResDur1 varieties. Indeed, no other cases of resistance breakdown in varieties combining *Rpv1* and *Rpv3.1* have been reported at the regional scale.

In this study, all *P. viticola* strains, except one, exhibited high sporulation ratios compared to avirulent reference strains on resistant cultivars carrying *Rpv1* alone or in combination with *Rpv3.1*. A similar result was observed for vir3.1 reference strains, despite their avirulent profile on *Rpv1*. These results suggest that sporulation levels on *Rpv1* cultivars are not correlated with the occurrence of a HR. The leaf disc assays conducted under controlled laboratory conditions may not fully reflect the reduction in sporulation associated with *Rpv1* resistance that is typically observed under field conditions (Calonnec *et al*., 2013). Nevertheless, our results clearly identify that at least four strains failed to induce any HR on the twelve leaf discs of varieties carrying *Rpv1*, indicating that these strains have acquired an additional virulence absent in the *Rpv3.1* reference strains. A gradient of virulence profiles toward *Rpv1* was observed, ranging from strains with complete adaptation (0/12 leaf discs showing HR) to almost complete avirulence (10/12 discs with HR) (**Figure S3**). Higher levels of adaptation to *Rpv1* had no detectable effect on sporulation in *Rpv1* varieties (Coutia, Artaban, Floreal), but were linked to reduced sporulation on Regent (*Rpv3.1*) (**Figure S5**). This observation suggests that acquiring a double virulence may compromise the ability of strains to sporulate effectively on hosts with a single resistance factor. Such a fitness trade-off is commonly encountered when pathogen develops multiple resistances to stress (Wang *et al*., 2017). This question remains to be explored in the case of multiple virulence against partial resistances.

Compared to avirulent strains, the four adapted strains show lower sporulation ratios on the susceptible variety Chardonnay (**Figure 3**). Pathogen adaptation may be penalized by a fitness cost on susceptible hosts, resulting in a decreased ability of resistance-adapted strains to infect susceptible hosts compared to non-adapted pathogens (Leach *et al*., 2001; Mundt, 2002, 2014; Thrall & Burdon, 2003). Modeling studies have shown that the existence of fitness costs is a major factor to improve the durability of both qualitative and quantitative resistances (Rimbaud *et al*., 2021). Yet, virulence costs have never been detected in the cases of resistance breakdown reported in *P. viticola* (Peressotti *et al*., 2010; Delmotte *et al*., 2014; Delmas *et al*., 2016; Heyman *et al*., 2021; Paineau *et al*., 2022). Interestingly, our results suggest that the virulence costs can be detectable on one susceptible cultivar (here Chardonnay) but not on another (Cabernet Sauvignon). It is well-established that grapevine cultivars exhibit different levels of susceptibility to downy mildew (Boso Alonso & Kassemeyer, 2008; Cadle-Davidson, 2008; Boso *et al*., 2011). Our results underscore the importance of evaluating potential adaptation costs across a range of representative susceptible cultivars.

Pyramiding can be an effective strategy to limit pathogen adaptation mainly as it decrease the probability that the pathogen crossed the mutational pathway conferring adaption to each resistance gene (McDonald & Linde, 2002; Rimbaud *et al*., 2021; Gandon *et al*., 2024). However, concurrently deploying a pyramided variety along with single-gene-resistant varieties, carrying the resistance genes stacked into the pyramid, result in a drastic decrease of the evolutionary and epidemiological control provided by pyramids (Brown, 2015; Horgan *et al*., 2018; Zaffaroni *et al*., 2024). The adaptation of *P. viticola* to *Rpv3.1* factor is long-standing following the early use of French-American hybrids carrying *Rpv3.1* since the early 20th century (Di Gaspero *et al*., 2012; Delmotte *et al*., 2014; Paineau *et al*., 2022), and likely widespread in European populations. The widespread presence of the vir3.1 alleles (>90% of the strains sampled in our study) represents a threat to the durability of resistance in varieties pyramiding *Rpv3.1* with other resistance factors. Indeed, several studies have already reported the existence of multi-virulent strains combining vir3.1 with either vir10 or vir12, confirming that pyramided varieties relying on *Rpv3.1* may be particularly vulnerable to breakdown. A similar scenario likely occurred in our study, leading to the breakdown of *Rpv1*.

In conclusion, our findings raise concern about the durability of the genetic basis of downy mildew resistance in commercially available varieties which primarily relies on *Rpv1*, *Rpv3.1*, *Rpv10* and *Rpv12* (Merdinoglu *et al*., 2018). Despite their recent and limited deployment in vineyards (representing approximately 0.36% of the french vineyard in 2024), cases of breakdowns for all four resistance factor have now been reported in the field, and confirmed in controlled conditions (Peressotti *et al*., 2010; Delmotte *et al*., 2014; Heyman *et al*., 2021; Wingerter *et al*., 2021; Paineau *et al*., 2022). Identifying the genomic determinant of *Rpv1* virulence, and assessing potential fitness costs associated with these mutations during the entire pathogen life cycle, should be considered priorities to guide breeding and deployment strategies. While further investigations are needed to determine whether *Rpv1-Rpv3.1* breaking strains have emerged in other wine growing regions, our results highlight the importance of monitoring virulence frequencies in the field, especially in regions where monogenic and pyramided resistances are deployed in close geographic proximity. This is particularly critical for perennial crops like grapevine, where resistance genes are intended to remain effective over extended periods (Zaffaroni *et al*., 2024). In practice, monitoring the emergence and spread of virulence within epidemiological surveillance networks (Soubeyrand *et al*., 2024) is indeed essential for ensuring sustainable and resilient cropping systems.

## References

Agurto M, Schlechter RO, Armijo G, Solano E, Serrano C, Contreras RA, Zúñiga GE, Arce-Johnson P. 2017. RUN1 and REN1 Pyramiding in Grapevine (Vitis vinifera cv. Crimson Seedless) Displays an Improved Defense Response Leading to Enhanced Resistance to Powdery Mildew (Erysiphe necator). Frontiers in Plant Science 8.

Bates D, Mächler M, Bolker B, Walker S. 2015. Fitting Linear Mixed-Effects Models Using lme4. Journal of Statistical Software 67: 1–48.

Bilan climatique 2024 en France | Météo-France.

Boso Alonso S, Kassemeyer HH. 2008. Different susceptibility of European grapevine cultivars for downy mildew.

Boso S, Alonso-Villaverde V, Gago P, Santiago J l., Martínez M c. 2011. Susceptibility of 44 grapevine (Vitis vinifera L.) varieties to downy mildew in the field. Australian Journal of Grape and Wine Research 17: 394–400.

Bove F, Bavaresco L, Caffi T, Rossi V. 2019. Assessment of Resistance Components for Improved Phenotyping of Grapevine Varieties Resistant to Downy Mildew. Frontiers in Plant Science 10.

Bretz F, Hothorn T, Westfall P. 2008. Multiple Comparison Procedures in Linear Models. In: Brito P, ed. COMPSTAT 2008. Heidelberg: Physica-Verlag HD, 423–431.

Brown JKM. 2015. Durable Resistance of Crops to Disease: A Darwinian Perspective. Annual Review of Phytopathology 53: 513–539.

Bruvo R, Michiels NK, D’souza TG, Schulenburg H. 2004. A simple method for the calculation of microsatellite genotype distances irrespective of ploidy level. Molecular Ecology 13: 2101–2106.

Cadle-Davidson L. 2008. Variation Within and Between Vitis spp. for Foliar Resistance to the Downy Mildew Pathogen Plasmopara viticola. Plant Disease 92: 1577–1584.

Calonnec A, Wiedemann-Merdinoglu S, Delière L, Cartolaro P, Schneider C, Delmotte F. 2013. The reliability of leaf bioassays for predicting disease resistance on fruit: a case study on grapevine resistance to downy and powdery mildew. Plant Pathology 62: 533–544.

Delière L, Cartolaro P, Léger B, Naud O. 2015. Field evaluation of an expertise-based formal decision system for fungicide management of grapevine downy and powdery mildews. Pest Management Science 71: 1247–1257.

Delmas CEL, Fabre F, Jolivet J, Mazet ID, Richart Cervera S, Delière L, Delmotte F. 2016. Adaptation of a plant pathogen to partial host resistance: selection for greater aggressiveness in grapevine downy mildew. Evolutionary Applications 9: 709–725.

Delmotte F, Chen WJ, Richard-Cervera S, Greif C, Papura D, Giresse X, Mondor-Genson G, Corio-Costet MF. 2006. Microsatellite DNA markers for Plasmopara viticola, the causal agent of downy mildew of grapes. Molecular Ecology Notes 6: 379–381.

Delmotte F, Mestre P, Schneider C, Kassemeyer H-H, Kozma P, Richart-Cervera S, Rouxel M, Delière L. 2014. Rapid and multiregional adaptation to host partial resistance in a plant pathogenic oomycete: Evidence from European populations of *Plasmopara viticola*, the causal agent of grapevine downy mildew. Infection, Genetics and Evolution 27: 500–508.

Di Gaspero G, Copetti D, Coleman C, Castellarin SD, Eibach R, Kozma P, Lacombe T, Gambetta G, Zvyagin A, Cindrić P, et al. 2012. Selective sweep at the Rpv3 locus during grapevine breeding for downy mildew resistance. Theoretical and Applied Genetics 124: 277–286.

Dussert Y, Mazet ID, Couture C, Gouzy J, Piron M-C, Kuchly C, Bouchez O, Rispe C, Mestre P, Delmotte F. 2019. A High-Quality Grapevine Downy Mildew Genome Assembly Reveals Rapidly Evolving and Lineage-Specific Putative Host Adaptation Genes. Genome Biology and Evolution 11: 954–969.

Dvorak E, Dumartinet T, Mazet ID, Chataigner A, Paineau M, Cantù D, Mestre P, Foulongne-Oriol M, Delmotte F. 2025. Parallel adaptation and admixture drive the evolution of virulence in the grapevine downy mildew pathogen.: 2025.05.18.654733.

Feechan A, Anderson C, Torregrosa L, Jermakow A, Mestre P, Wiedemann-Merdinoglu S, Merdinoglu D, Walker AR, Cadle-Davidson L, Reisch B, et al. 2013. Genetic dissection of a TIR-NB-LRR locus from the wild North American grapevine species Muscadinia rotundifolia identifies paralogous genes conferring resistance to major fungal and oomycete pathogens in cultivated grapevine. The Plant Journal 76: 661–674.

Fontaine MC, Labbé F, Dussert Y, Delière L, Richart-Cervera S, Giraud T, Delmotte F. 2021. Europe as a bridgehead in the worldwide invasion history of grapevine downy mildew, Plasmopara viticola. Current Biology 31: 2155–2166.e4.

Foria S, Magris G, Morgante M, Di Gaspero G. 2018. The genetic background modulates the intensity of Rpv3-dependent downy mildew resistance in grapevine. Plant Breeding 137: 220–228.

Gandon S, Guillemet M, Gatchitch F, Nicot A, Renaud AC, Tremblay DM, Moineau S. 2024. Building pyramids against the evolutionary emergence of pathogens. Proceedings of the Royal Society B: Biological Sciences 291: 20231529.

Gessler C, Pertot I, Perazzolli M. 2011. Plasmopara viticola: a review of knowledge on downy mildew of grapevine and effective disease management. Phytopathologia Mediterranea 50: 3–44.

Guimier S, Delmotte F, Miclot AS, Fabre F, Mazet I, Couture C, Schneider C, Delière L. 2019. OSCAR, a national observatory to support the durable deployment of disease-resistant grapevine cultivars. Acta Horticulturae: 21–34.

Heyman L, Höfle R, Kicherer A, Trapp O, Ait Barka E, Töpfer R, Höfte M. 2021. The Durability of Quantitative Host Resistance and Variability in Pathogen Virulence in the Interaction Between European Grapevine Cultivars and Plasmopara viticola. Frontiers in Agronomy 3.

Horgan FG, Bernal CC, Vu Q, Almazan MLP, Ramal AF, Yasui H, Fujita D. 2018. Virulence adaptation in a rice leafhopper: Exposure to ineffective genes compromises pyramided resistance. Crop Protection 113: 40–47.

Kamvar ZN, Tabima JF, Grünwald NJ. 2014. Poppr: an R package for genetic analysis of populations with clonal, partially clonal, and/or sexual reproduction. PeerJ 2: e281.

Koledenkova K, Esmaeel Q, Jacquard C, Nowak J, Clément C, Ait Barka E. 2022. Plasmopara viticola the Causal Agent of Downy Mildew of Grapevine: From Its Taxonomy to Disease Management. Frontiers in Microbiology 13.

Leach JE, Cruz CMV, Bai J, Leung H. 2001. PATHOGEN FITNESS PENALTY AS A PREDICTOR OF DURABILITY OF DISEASE RESISTANCE GENES. Annual Review of Phytopathology 39: 187–224.

Lenth RV. 2017. emmeans: Estimated Marginal Means, aka Least-Squares Means.

McDonald BA, Linde C. 2002. PATHOGEN POPULATION GENETICS, EVOLUTIONARY POTENTIAL, AND DURABLE RESISTANCE. Annual Review of Phytopathology 40: 349–379.

Merdinoglu D, Schneider C, Prado E, Wiedemann-Merdinoglu S, Mestre P. 2018. Breeding for durable resistance to downy and powdery mildew in grapevine. OENO One 52: 203–209.

Merdinoglu D, Wiedeman-Merdinoglu S, Coste P, Dumas V, Haetty S, Butterlin G, Greif C. 2003. GENETIC ANALYSIS OF DOWNY MILDEW RESISTANCE DERIVED FROM MUSCADINIA ROTUNDIFOLIA. Acta Horticulturae: 451–456.

Miclot AS, Delmotte F, Bourg J, Mazet ID, Fabre F, Delière L. 2022. Four years of monitoring of disease-resistant grapevine varieties in French vineyards (T Caffi, V Rossi, and G Fedele, Eds.). BIO Web of Conferences 50: 02008.

Millardet A. 1881. Notes sur les vignes américaines et opuscules divers sur le même sujet. Feret.

Möller EM, Bahnweg G, Sandermann H, Geiger HH. 1992. A simple and efficient protocol for isolation of high molecular weight DNA from filamentous fungi, fruit bodies, and infected plant tissues. Nucleic Acids Research 20: 6115–6116.

Mundt CC. 2002. Use of multiline cultivars and cultivar mixtures for disease management. Annual Review of Phytopathology 40: 381–410.

Mundt CC. 2014. Durable resistance: A key to sustainable management of pathogens and pests. *Infection*, Genetics and Evolution 27: 446–455.

Nefti O, Chartier N, Merot A, Peyrard T, Delière L. 2024. To what extent can a phase-out of pesticides in viticulture be achieved? Learning from the efforts of a large farm network after 10 years. OENO One 58.

Paineau M, Mazet ID, Wiedemann-Merdinoglu S, Fabre F, Delmotte F. 2022. The Characterization of Pathotypes in Grapevine Downy Mildew Provides Insights into the Breakdown of Rpv3, Rpv10, and Rpv12 Factors in Grapevines. Phytopathology® 112: 2329–2340.

Paineau M, Minio A, Mestre P, Fabre F, Mazet ID, Couture C, Legeai F, Dumartinet T, Cantu D, Delmotte F. 2024. Multiple deletions of candidate effector genes lead to the breakdown of partial grapevine resistance to downy mildew. New Phytologist 243: 1490–1505.

Paradis E, Claude J, Strimmer K. 2004. APE: Analyses of Phylogenetics and Evolution in R language. *Bioinformatics (Oxford*, England) 20: 289–290.

Parlevliet JE. 2002. Durability of resistance against fungal, bacterial and viral pathogens; present situation. Euphytica 124: 147–156.

Peressotti E, Wiedemann-Merdinoglu S, Delmotte F, Bellin D, Di Gaspero G, Testolin R, Merdinoglu D, Mestre P. 2010. Breakdown of resistance to grapevine downy mildew upon limited deployment of a resistant variety. BMC Plant Biology 10: 147.

Possamai T, Wiedemann-Merdinoglu S. 2022. Phenotyping for QTL identification: A case study of resistance to Plasmopara viticola and Erysiphe necator in grapevine. Frontiers in Plant Science 13.

Rimbaud L, Fabre F, Papaïx J, Moury B, Lannou C, Barrett LG, Thrall PH. 2021. Models of Plant Resistance Deployment. Annual Review of Phytopathology 59: 125–152.

Rouxel M, Papura D, Nogueira M, Machefer V, Dezette D, Richard-Cervera S, Carrere S, Mestre P, Delmotte F. 2012. Microsatellite Markers for Characterization of Native and Introduced Populations of Plasmopara viticola, the Causal Agent of Grapevine Downy Mildew. Applied and Environmental Microbiology 78: 6337–6340.

Salmon J-M, Ojeda H, Escudier J-L. 2018. Disease resistant grapevine varieties and quality: the case of Bouquet varieties. OENO One 52: 6 p.

Schneider C, Onimus C, Prado E, Dumas V, Wiedemann-Merdinoglu S, Dorne MA, Lacombe MC, Piron MC, Umar-Faruk A, Duchêne E, et al. 2019. INRA-ResDur: the French grapevine breeding programme for durable resistance to downy and powdery mildew. Acta Horticulturae: 207–214.

Soubeyrand S, Estoup A, Cruaud A, Malembic-Maher S, Meynard C, Ravigné V, Barbier M, Barrès B, Berthier K, Boitard S, et al. 2024. Building integrated plant health surveillance: a proactive research agenda for anticipating and mitigating disease and pest emergence. CABI Agriculture and Bioscience 5: 72.

Thind TS, Arora JK, Mohan C, Raj P. 2004. Epidemiology of Powdery Mildew, Downy Mildew and Anthracnose Diseases of Grapevine. In: Naqvi SAMH, ed. Diseases of Fruits and Vegetables Volume I: Diagnosis and Management. Dordrecht: Springer Netherlands, 621–638.

Thrall PH, Burdon JJ. 2003. Evolution of Virulence in a Plant Host-Pathogen Metapopulation. Science 299: 1735–1737.

Wang X, Wei Z, Li M, Wang X, Shan A, Mei X, Jousset A, Shen Q, Xu Y, Friman V-P. 2017. Parasites and competitors suppress bacterial pathogen synergistically due to evolutionary trade-offs. Evolution; International Journal of Organic Evolution 71: 733–746.

Wickham H. 2016. ggplot2. Cham: Springer International Publishing.

Wingerter C, Eisenmann B, Weber P, Dry I, Bogs J. 2021. Grapevine Rpv3-, Rpv10-and Rpv12-mediated defense responses against Plasmopara viticola and the impact of their deployment on fungicide use in viticulture. BMC Plant Biology 21: 470.

Yobrégat O. 2018. Introduction to resistant vine types: a brief history and overview of the situation. OENO One 52: 241–246.

Yu G, Smith DK, Zhu H, Guan Y, Lam TT-Y. 2017. ggtree: an r package for visualization and annotation of phylogenetic trees with their covariates and other associated data. Methods in Ecology and Evolution 8: 28–36.

Zaffaroni M, Papaïx J, Geffersa AG, Rey J-F, Rimbaud L, Fabre F. 2024. Combining Single-Gene-Resistant and Pyramided Cultivars of Perennial Crops in Agricultural Landscapes Compromises Pyramiding Benefits in Most Production Situations. Phytopathology® 114: 2310–2321.

