## supplementary material for "Multiple geographic breakdown events of the *Rpv1*–*Rpv3.1* pyramided resistance in grapevine by *Plasmopara viticola*"

### Supplementary data

**Table S1: Virulence profiles and origin of the 8 reference strains of *Plasmopara viticola* used for inoculation**

| Virulence profile | Strain ID | Source host | QTL in source host | Year of collection | Country | Locality | Reference |
|---|---|---|---|---|---|---|---|
| vir3.1 | PV2664_1 | Artaban | *Rpv1*, *Rpv3.1* | 2013 | France | Villenave d'Ornon | Paineau et al. (2022) |
| vir3.1 | PV412_11 | Regent | *Rpv*3.1 | 2010 | Switzerland | Cugnasco | Paineau et al. (2022) |
| vir3.1,12 | PV8522_1 | Unknown | *Rpv*3.1, *Rpv12* | NA | Switzerland | Soyhières | Wingerter et al. (2021) |
| avir | PV221_2 | Sauvignon | no resistance | 2009 | France | Blanquefort | Paineau et al. (2024) |
| avir | PV2578_1 | Chardonnay | no resistance | 2013 | France | Le Landreau | Paineau et al. (2022) |
| avir | PV2598_11 | Chardonnay | no resistance | 2013 | Spain | Agoncillo | Paineau et al. (2022) |
| avir | PV2909_11 | Chardonnay | no resistance | 2014 | Germany | Geisenheim | Paineau et al. (2022) |
| avir | PV3174_11 | Nebbiolo | no resistance | 2016 | Italy | Piateda | Paineau et al. (2022) |

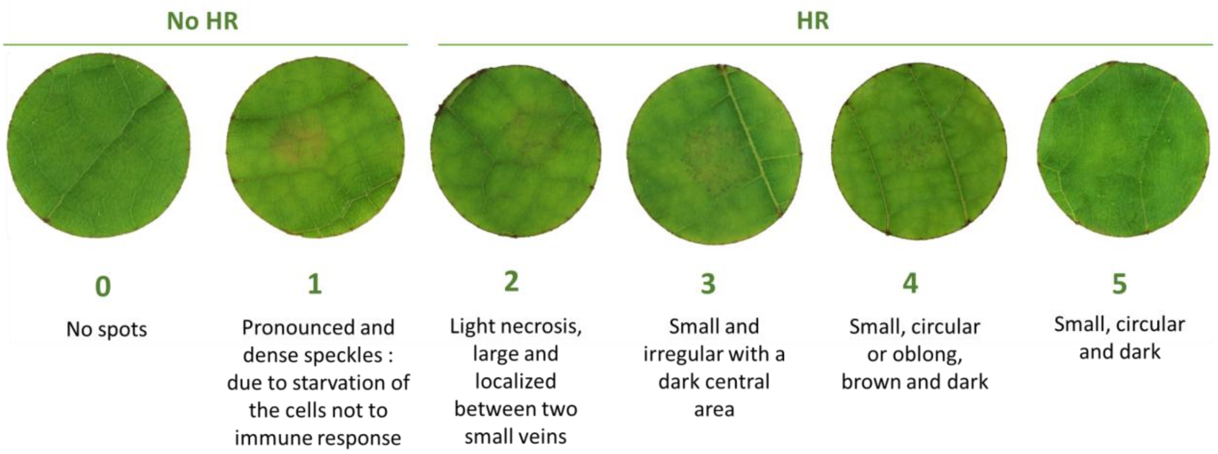


**Figure S1. Necrosis types scale.**

The qualitative necrosis scale used to assess Hypersensitive Response (HR) to *P. viticola* infection on grapevine leaf discs is adapted from Paineau et al. 2022 (see FigS1). On the top of the figure is displayed the classification of the necrosis types corresponding to the absence (no HR) or presence of HR (HR). Note that the score did not correspond to an ordinal score of necrosis but characterize the type of necrosis as a qualitative variable.

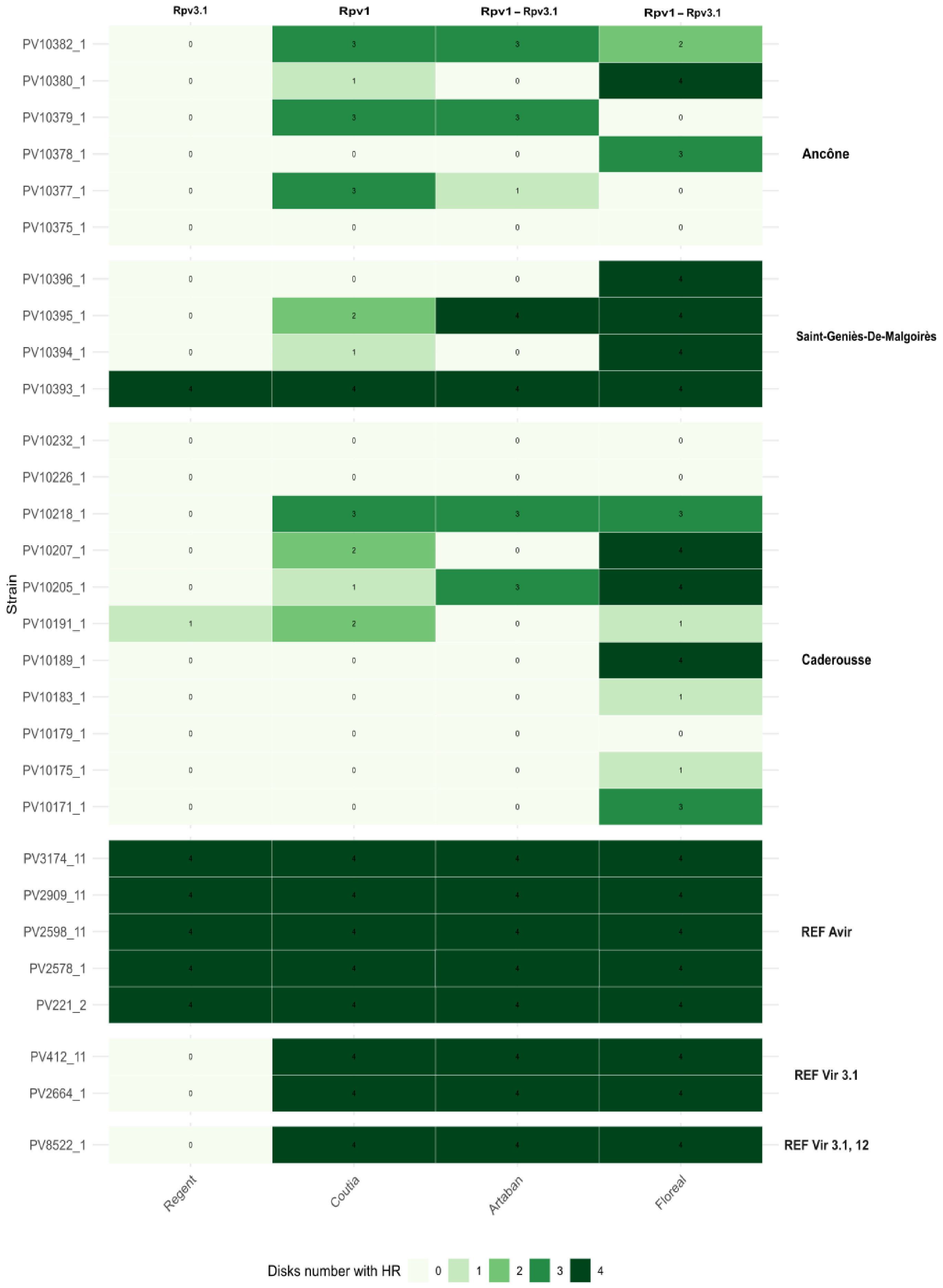
Rpv3.1
Rpv1
Rpv1 – Rpv3.1
Rpv1 – Rpv3.1
Strain
PV10382_1
PV10380_1
PV10379_1
PV10378_1
PV10377_1
PV10375_1
Ancône
PV10396_1
PV10395_1
PV10394_1
PV10393_1
Saint-Geniès-De-Malgoirès
PV10232_1
PV10226_1
PV10218_1
PV10207_1
PV10205_1
PV10191_1
PV10189_1
PV10183_1
PV10179_1
PV10175_1
PV10171_1
Caderousse
PV3174_11
PV2909_11
PV2598_11
PV2578_1
PV221_2
REF Avir
PV412_11
PV2664_1
REF Vir 3.1
PV8522_1
REF Vir 3.1, 12
Regent
Coutia
Artaban
Floreal
Disks number with HR
0
1
2
3
4

**Figure S2. Hypersensitive response profiles of the 21 downy mildew strains collected in the field and the 8 reference strains.**

Number of leaf discs (out of 4 replicates) with necrosis scores from 2 to 5 (HR) on grapevine resistant varieties: Regent (*Rpv3.1*), Coutia (*Rpv1*), Artaban and Floreal (*Rpv1-Rpv3.1*) are represented. Strains origin is shown on the right: avirulent, virulent references *vir3.1* and *vir3.1,12*, and the field strains from the 3 different location. Strains with 0/4 discs showing HR are classified as virulent on the variety tested, whereas strains presenting at least one disc with necrosis are classified as avirulent on the variety tested.

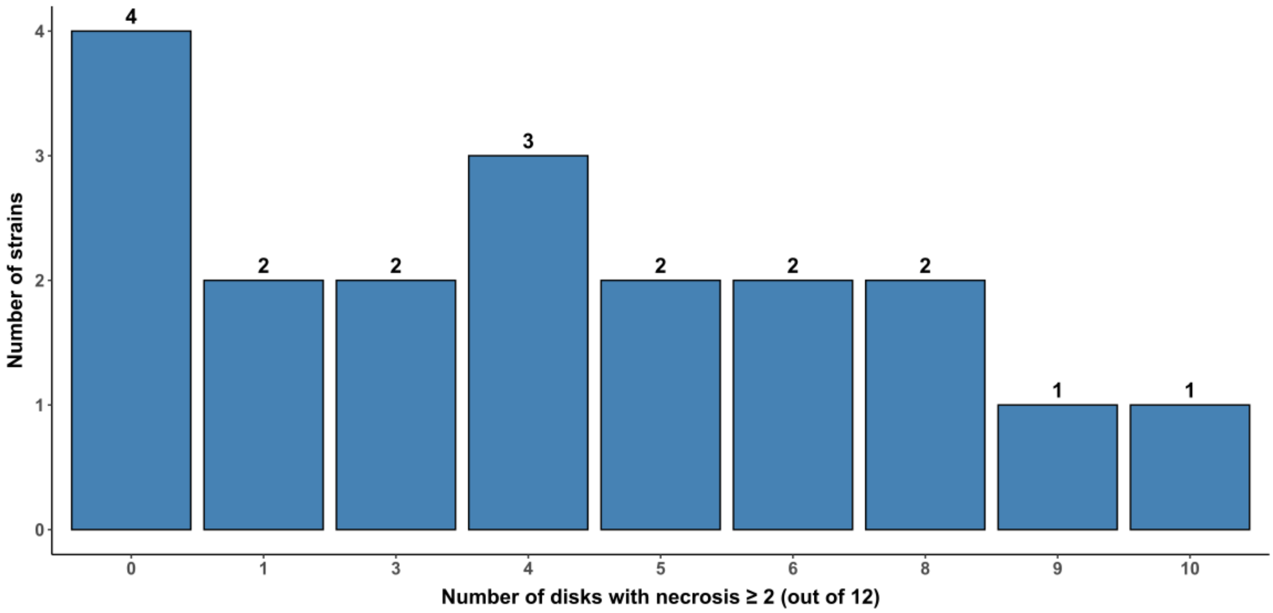


**Figure S3. Distribution of virulence profiles of strains regarding the *Rpv1* factor.**

Distribution of the number of leaf discs with necrosis scores ≥2 (HR) on grapevine varieties carrying the resistance factor *Rpv1* alone or pyramided with *Rpv3.1*. Only the 19 field strains previously identified as *vir3.1* (i.e., no discs with HR on the *Rpv3.1*-only variety Regent) are shown. Strains are ranked from fully adapted (0/12 HR discs) to non-adapted (up to 10/12 HR discs), based on symptoms observed on Coutia (*Rpv1*), Artaban, and Floreal (*Rpv1*-*Rpv3.1*) (Four discs per variety, resulting in a total of 12 discs (4 × 3)

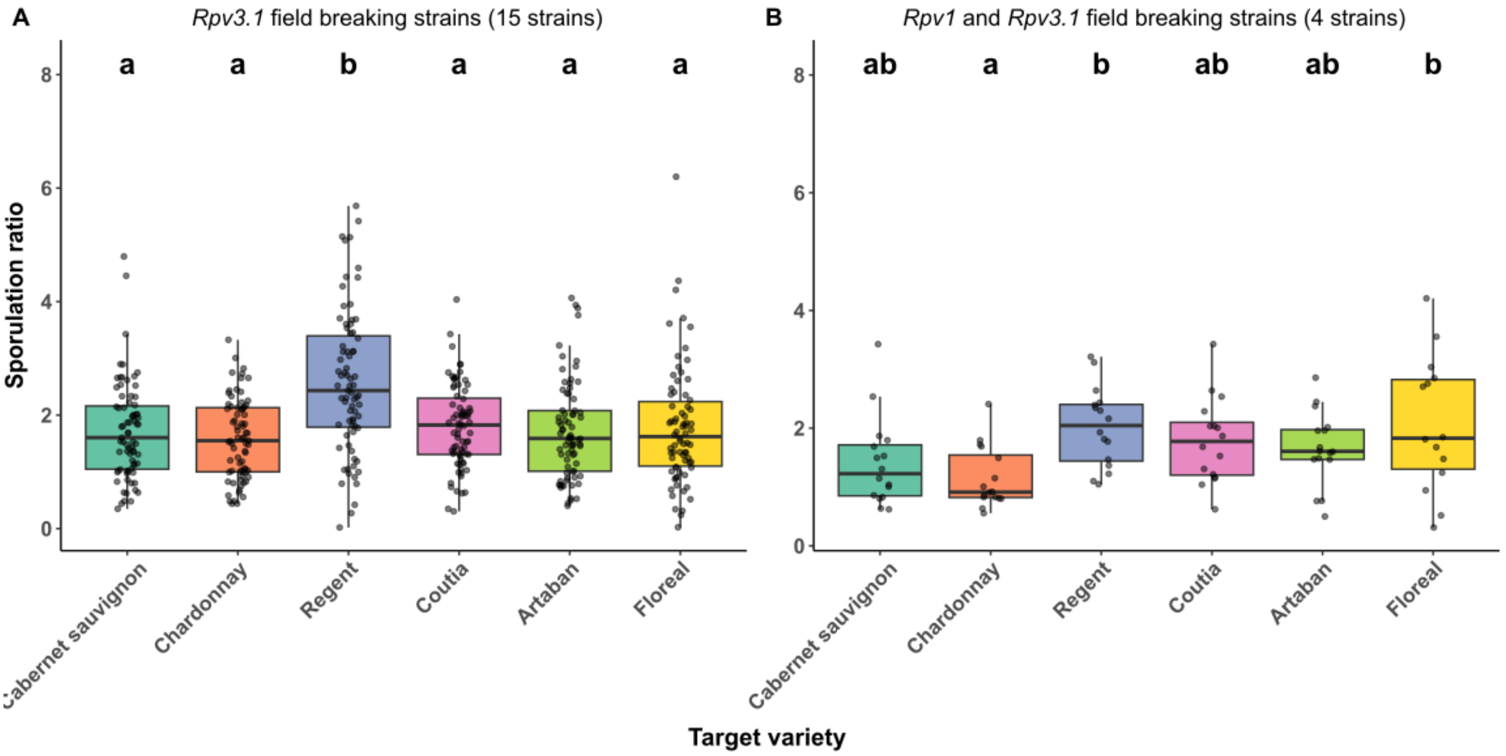


**Figure S4. Comparison of sporulation ratio of the *Rpv3.1* breaking strains and the four *Rpv1, Rpv3.1* breaking strains.**

Boxplots show the sporulation ratio (expressed as a percentage of sporulating leaf disc surface) for A) the 15 strains identified as only *Rpv3.1* breaking strains (n=60 for each boxplot) and B) for the 4 *Rpv1-Rpv3.1* breaking strains (n=16 for each boxplot), observed on the six grapevine varieties inoculated: Chardonnay and Cabernet Sauvignon (susceptible), Coutia (*Rpv1*), Regent (*Rpv3.1*), Artaban and Floreal (*Rpv1-Rpv3.1*). Sporulation ratio was assessed via image analysis. Letters above the boxplots indicate significant groupings based on Tukey's HSD post-hoc test ($\alpha = 0.05$).

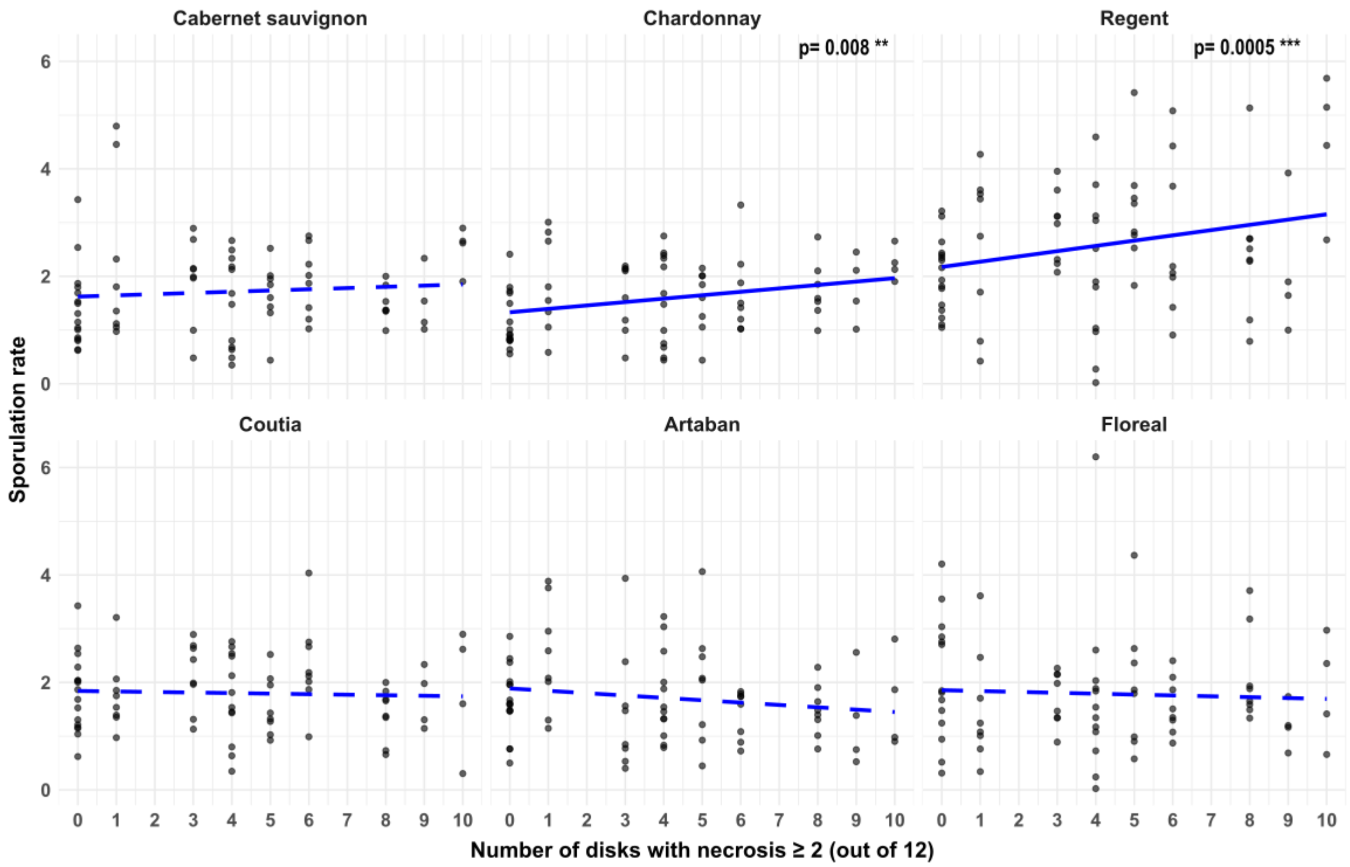


**Figure S5. Relationship between adaptation to *Rpv1* factor and sporulation ratio across varieties.**

Sporulation ratio is represented for 19 field strains previously identified as *Rpv3.1* breaking (*vir3.1*), based on the absence of HR on Regent (*Rpv3.1*), as described in methods and **Figure.S3**. Strains are grouped according to the number of necrotic discs (score ≥ 2) observed on three varieties carrying the *Rpv1* factor: Coutia (*Rpv1*), Artaban (*Rpv1-Rpv3.1*) and Floreal (*Rpv1-Rpv3.1*) out of the 12 discs inoculated (4 discs per variety). The number of necrotic discs reflect a gradient of adaptation to the *Rpv1* resistance factor, from fully adapted strains (0/12 necrotic discs) to poorly adapted strains (up to 10/12). The blue line represents the regression line between sporulation ratio and adaptation level (number of necrotic discs) as fitted with a linear model including an interaction between the inoculated variety and the number of necrotic. A solid line represents a significative interaction.
